# Cell Numbers Contribute to Cell Fate During Ciona Cardiopharyngeal Mesoderm Specification

**DOI:** 10.1101/2025.11.10.687625

**Authors:** Emily Singer, Haram Kim, Michael Levine, Nicholas Treen

## Abstract

The *Ciona* heart cell lineage can be accurately traced back to a pair of blastomeres, the B7.5 cells, that form at the 64-cell stage. In addition to the adult heart, the B7.5 cells also contribute to two tail muscle cells in the larva, as well as the muscles that form the siphons for pumping water for feeding. Because of the simplicity of this system, we have a good understanding of how the B7.5 derivatives are specified during development. However, we know less about how the *Ciona* embryo precisely regulates cell numbers, as well as what effects altering cell numbers will have on development. We found that cell numbers in the B7.5 lineage are controlled by a pulse of transcription of the cell cycle inhibitor *Cdkn1.b*. *Cdkn1.b* can be repressed by the paralogues *Prdm1-r.a* and *Prdm1-r.b* that are exclusively transcribed in the B7.5 cells at the 110-cell stage. We unexpectedly found that precocious arrest of cell division in the B7.5 cell lineage resulted in a reversion to tail muscle fate, even in cells that can migrate. Our work demonstrates an unexpected connection between the control of cell numbers and cell fate in development.

**Significance Statement:** The control of cell numbers is essential to many developmental processes, but it has been relatively understudied. A clear example of this can be seen in heart development. Hearts develop from precursor cells called cardiopharyngeal mesoderm. These cells must be able to migrate and proliferate in a precise way or heart defects will occur. We took advantage of the simplicity of embryonic development in the chordate *Ciona* (an invertebrate closely related to vertebrates) to investigate the consequences of disrupting cell numbers on heart development. Our finding that an early arrest in cell division results in a failure of heart development has implications for similar processes in vertebrates, including humans.

## Introduction

The control of cell numbers during embryogenesis is kept under strict control since the presence of too many or too few cells can result in developmental defects (1). Despite its central importance, we have a limited understanding of how cell numbers are controlled and the consequences of altering these numbers. A major limiting factor is the large number of cells comprising most embryos, especially vertebrate embryos, which often contain thousands to millions of cells.

Ascidians such as *Ciona* are well suited for analyzing the control of cell division in development as their embryos consist of a remarkably low number of cells, less than 2000 at the time of hatching (2). Ascidians are also the invertebrates most closely related to vertebrates and possess rudiments of several defining vertebrate features such as neural crest (3) and cardiopharyngeal mesoderm (4), allowing findings in the simple *Ciona* embryo to be easily recontextualized in vertebrates.

The vertebrate cardiopharyngeal mesoderm is an embryonic territory that produces the heart as well as branchiomeric muscles in the throat and neck (5). The cardiopharyngeal mesoderm may have been at least as important as the neural crest in the evolution of the vertebrate “new head” (6). Ascidians have a clear equivalent of the vertebrate cardiopharyngeal mesoderm, arising from trunk ventral cells (TVCs) (7). The *Ciona* TVCs/cardiopharyngeal mesoderm produce the adult heart, as well as the muscles in the atrial siphons that are used for feeding (4,7). Parallels between the ascidian and vertebrate cardiopharyngeal mesoderm are clear, as detailed studies of the TVCs show evidence for the existence of both first and second heart fields in *Ciona* embryos, along with conserved regulatory networks (8).

The *Ciona* cardiopharyngeal mesoderm contains remarkably few cells. This permitted a thorough analysis of its early specification, initially through the maternal factors Macho-1 (Zic-r.a) and β-catenin/Tcf7. These maternal factors can activate transcription of Tbx6 genes and *Lhx3/4* (9,10). *Tbx6-r.b* and *Lhx3/4* are typically expressed in the muscle and endoderm lineages respectively. However, when both are simultaneously expressed in the same cell, they trigger the activation of *Mesp*, a key determinant of the cardiopharyngeal mesoderm (11). This process only happens in a single pair of cells, the B7.5 cells, that are formed at the 64-cell stage. During gastrulation these cells divide twice to produce four granddaughter B9.17-20 cells (12). The two anterior cells receive an FGF signal that induces *Fox.f* expression, which is required to initiate a coordinated migration away from the tail and into the trunk. After migration they resume proliferation to form cardiopharyngeal mesoderm (8). The two posterior cells that do not migrate and remain in the tail to form anterior tail muscles that are morphologically indistinguishable from other muscle cells.

We took advantage of recently developed quantitative multiplexed *in situ* hybridization methods to reassess the control of cell numbers and cell fate in the development of the *Ciona* B7.5 blastomeres. We found an unexpected role for the *Ciona* homologues of Prdm1 (Blimp-1 in vertebrates) in this process through regulation of the cell cycle inhibitor Cdkn1.b. We also present evidence that precocious arrest of cell divisions results in the transformation of TVC progenitors into muscles, despite their migration from tail to trunk. These studies suggest that the specification of B7.5 cardiopharyngeal mesoderm depends on the integration of cell fate determinants (e.g. Mesp) and the control of the cell cycle by Prdm1/Cdkn1.b.

## Results

### *Prdm1-r.a/b* are exclusively transcribed in the B7.5 cells at the 110-cell stage

The *Ciona* heart is composed of cells that originate from the B7.5 blastomeres, which form at the 64-cell stage (Fig. 1A, 13). Both B7.5 blastomeres form four granddaughter cells consisting of two anterior tail muscles and two TVCs (13). The TVCs migrate from the tail to the trunk where they will continue to proliferate and differentiate into the beating heart and atrial siphon muscles during metamorphosis. The basic helix-loop-helix transcription factor *Mesp,* the *Ciona* homologue of vertebrate proteins Mesp1 and Mesp2, has a well-studied role in *Ciona* cardiopharyngeal mesoderm development and is notable because it is exclusively expressed in the B7.5 cells at the 110-cell stage (13). Previous single-cell RNA-seq datasets (14) suggest that the transcriptional repressors *Prdm1-r.a* and *Prdm1-r.b* are also predominantly expressed in B7.5 cells at the 110-cell stage (Fig. 1B). Other developmental genes that are expressed in the B7.5 cells, such as *Tbx6-r.b* and *Lhx3/4*, are also detected in other cell lineages such as developing tail muscles and endoderm precursors (Fig 1B).

**Fig. 1.**
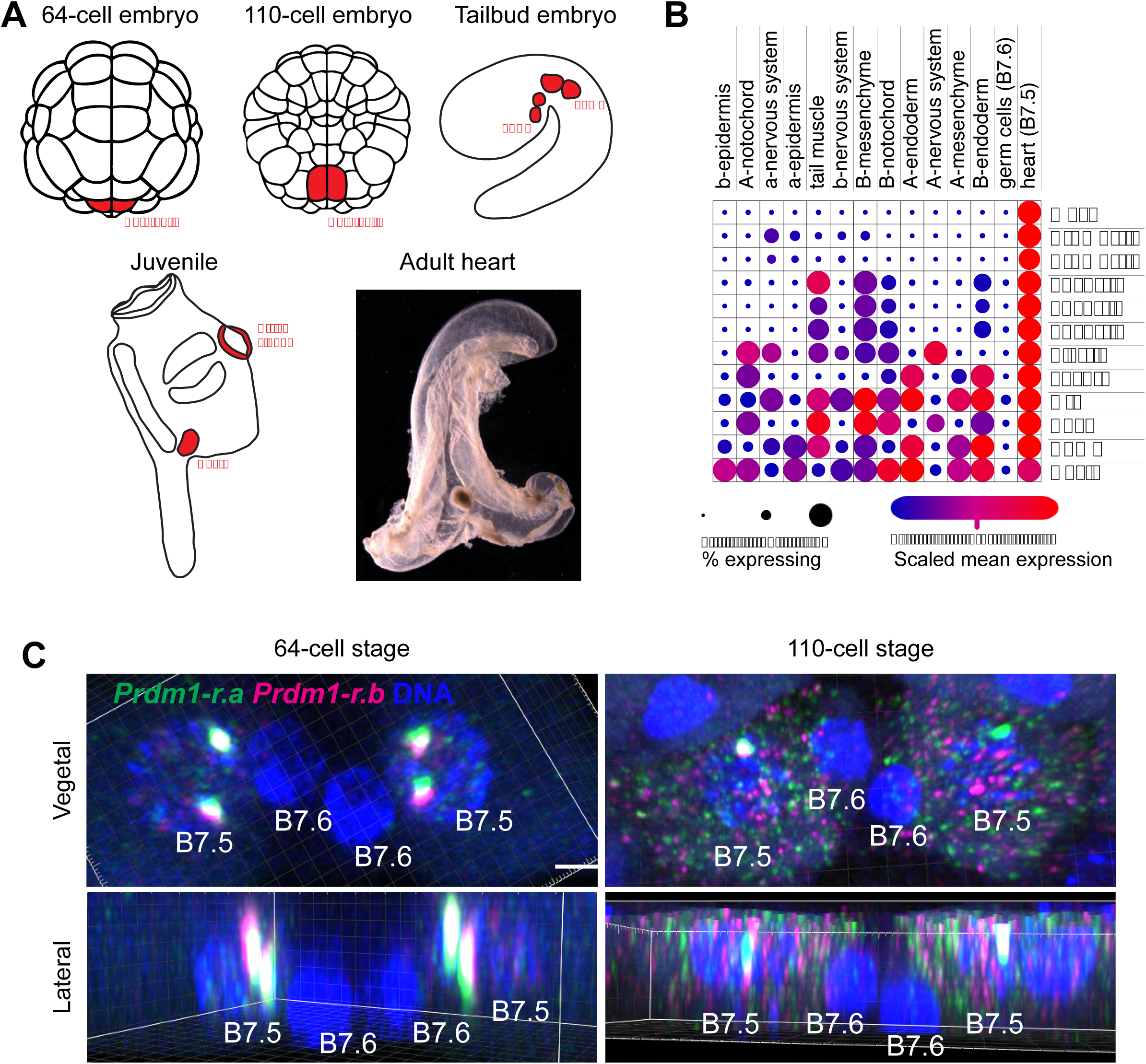
*Prdm1-r.a* and *Prdm1-r.b* are transcribed only in B7.5 cells at the 110-cell stage. (A) Schematic of *Ciona* embryos depicting the location of the B7.5 cells and their descendants. The 64-and 110-cell stage embryo depicts the cell outlines of the vegetal side of the embryo, anterior up. The tailbud embryo shows the granddaughter cells of a single B7.5 cell in one half of an embryo (anterior left, TVCs-trunk ventral cells, ATMs, anterior tail muscles). The juvenile shows the locations of the TVCs after metamorphosis in the heart and the atrial siphons. A photo is shown of an adult heart dissected from an individual *Ciona*. (B) Single-cell RNA-seq levels of *Prdm1-r.a*, *Prdm1-r.b*, as well as other genes that have previously described expression in *Ciona* embryos at the 110-cell stage. Cells are clustered into groups based on the predicted tissue type these cells will form later in development. (C) HCR *in-situ* hybridization signals for *Prdm1-r.a* and *Prdm1-r.b* in the B7.5 and B7.6 cells at the 64- and 110-cell stages. Images are 3D projections of confocal fluorescence microscopy datasets. Scale bar = 5 μm.

HCR *in situ* hybridization assays confirmed these observations. Most notably, *Prdm1-r.a* and *Prdm1-r.b* transcription occurs in the B7.5 cells at the 64-cell stage, but not in the transcriptionally silenced, sister cells (B7.6) that form germ cell precursors (Fig 1B,C). By the 110-cell stage, mRNAs for *Prdm1-r.a* and *Prdm1-r.b* are detected in B7.5 cytoplasm (Fig. 1C, S1). At the 64-cell stage there is some expression of *Prdm1-r.a* and *Prdm1-r.b* in the a7.13 cells (Fig. S1, 15), but by the 110-cell stage they can only be reliably detected in the B7.5 cells (Fig. S1). This expression pattern suggested to us that *Prdm1-r.a* and *Prdm1-r.b* might have an unknown role in *Ciona* cardiopharyngeal mesoderm development.

### Timing of transcriptional activation in B7.5 cells

We systematically measured the level of active transcription for *Prdm1-r.a* and *Prdm1-r.b*, along with *Mesp* and other regulatory factors, in precisely staged embryos over a one-hour period at 7.5-minute intervals (Fig. 2). This period lasted from 3 hours 50 minutes post fertilization, when the B6.3 cell is in metaphase, throughout the 64-and 110-cell stages until the onset of gastrulation at 4 hours and 50 minutes post fertilization. Since the B6.3 precursor cells are transcriptionally silenced by the maternally segregated germline Pem1 repressor (16), these observations provide a readout of the transcriptional kinetics for zygotic genome activation in the B7.5 cells. Using 4-color HCR *in situ* hybridization we were able to recapitulate previously described gene expression patterns such as the expression of *Lhx3/4* in endoderm precursor cells, the expression of *Tbx6-r.b* in muscle precursors, and the expression of *Mesp* in B7.5, the only region of the embryo where *Lhx3/4* and *Tbx6-r.b* expression overlap (Fig. S2, 11). At 15 minutes following metaphase, *Lhx3/4*, *Tbx6-r.b/c/d*, and *Zic-r.b* all exhibit robust signals of active transcription in B7.5 nuclei prior to the onset of *Mesp* transcription (Fig. 2, S3). This situation persists until 30-40 minutes after metaphase when *Mesp* transcription first becomes reliably detectable (Fig. 2). Once *Mesp* transcription is active, the other genes we measured begin to decline and are mostly undetectable 60 minutes after metaphase, whereas *Mesp* transcription persists at high levels.

**Fig. 2.**
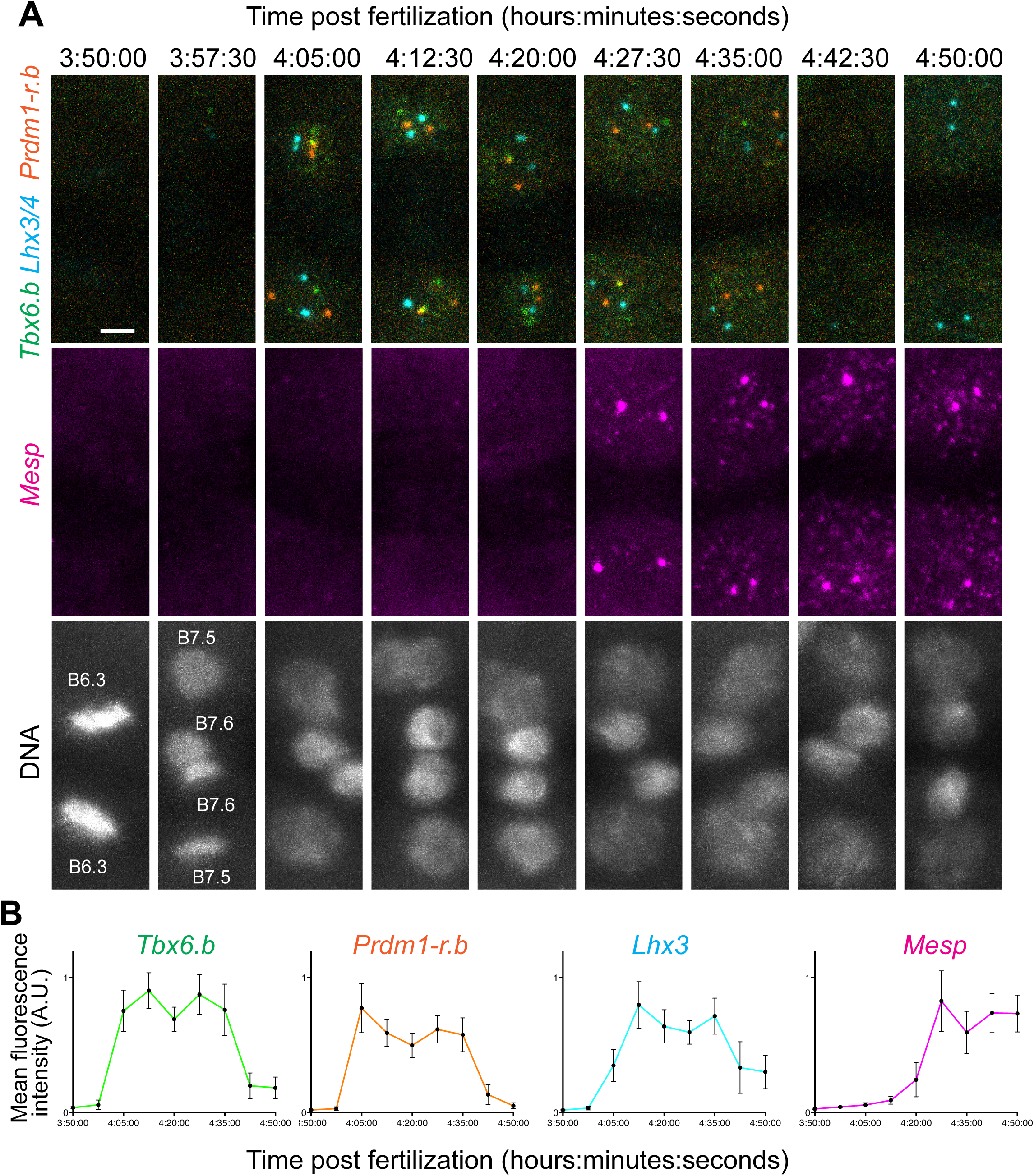
Onset of transcription in B7.5 Cells. (A) HCR *in situ* hybridization signals for *Tbx6-r.b*, *Lhx3/4*, *Prdm1-r.b*, and *Mesp* from the onset of mitosis of the B6.3 cells for 1 hour. Images are taken from individual fixed samples. The 3 images in a single time point are from the same embryo. Scale bar = 5 μm. (B) Quantification of levels of transcription from HCR *in-situ* hybridization signals for *Tbx6-r.b, Lhx3/4, Prdm1-r.b*, and *Mesp*. Each time point consists of 28 measurements from 7 embryos. Error bars plot 95% confidence intervals.

Transcriptional bursting has been observed in a variety of systems (17). In this regard we note general absence of obvious bursting in early *Ciona* embryos (18). Both copies of a gene were typically either on and transcribing at strong levels, or off with no detectable transcription. Transcription of a single copy of a gene could only be seen at the very start or end of interphase, and this state was probably extremely brief (18). Interestingly, *Lhx3/4* exhibits a wide range of variability in transcription in B7.5 cells at later time points (Fig. 2, S4) with transcription being off in around half the nuclei, as well as several examples of active transcription at single loci. These observations are consistent with *Lhx3/4* displaying transcriptional bursting after specification of defined cell lineages.

### *Prdm1-r.a* and *Prdm1-r.b* regulate B7.5 cell numbers

Co-transcription of *Prdm1-r.a* and *Prdm1-r.b* with known TVC determinants such as *Lhx3/4* and *Tbx6-r.b* suggests that Prdm1-r.a and *Prdm1-r.b* may have important roles in *Ciona* cardiopharyngeal mesoderm development. *Prdm1-r.a* and *Prdm1-r.b* are known to regulate *Zic-r.b* (15,19), which in turn, controls cell numbers in muscle, neural and notochord lineages by activating the expression of the cell cycle inhibitor *Cdkn1.b* (20,21). Because of these connections, we next investigated a role for *Prdm1-r.a* and *Prdm1-r.b* in controlling cell numbers in the B7.5 lineage.

The 5’ *Mesp* regulatory region was used to direct restricted expression of *Prdm1-r.a* and *Prdm1-r.b* in the B7.5 lineage (4). A *Mesp* reporter plasmid labeled all four granddaughter cells of each B7.5 cell at the tailbud stage (Fig. 3A). The anterior two cells were specifically labelled using a reporter plasmid containing the TVC enhancer from *Fox.f* (22, Fig. 3A). When *Cdkn1.b* is precociously expressed in the B7.5 cells there is only one cell division, resulting in just two daughter cells from each B7.5 blastomere (23, Fig 3. A.), both daughter cells express *Fox.f* and faithfully migrate at the expense of anterior tail muscles (Fig. 3A). When either *Prdm1-r.a* or *Prdm1-r.b* were overexpressed in B7.5, we observed an increase in cell numbers. The exact numbers were unpredictable and varied between replicates, but typically 8-12 cells were obtained from each B7.5 blastomere by the tailbud stage, suggesting three or four divisions rather than two. These cells fail to express the *Fox.f* reporter, suggesting that they were not specified to become TVCs and cannot migrate.

**Fig. 3.**
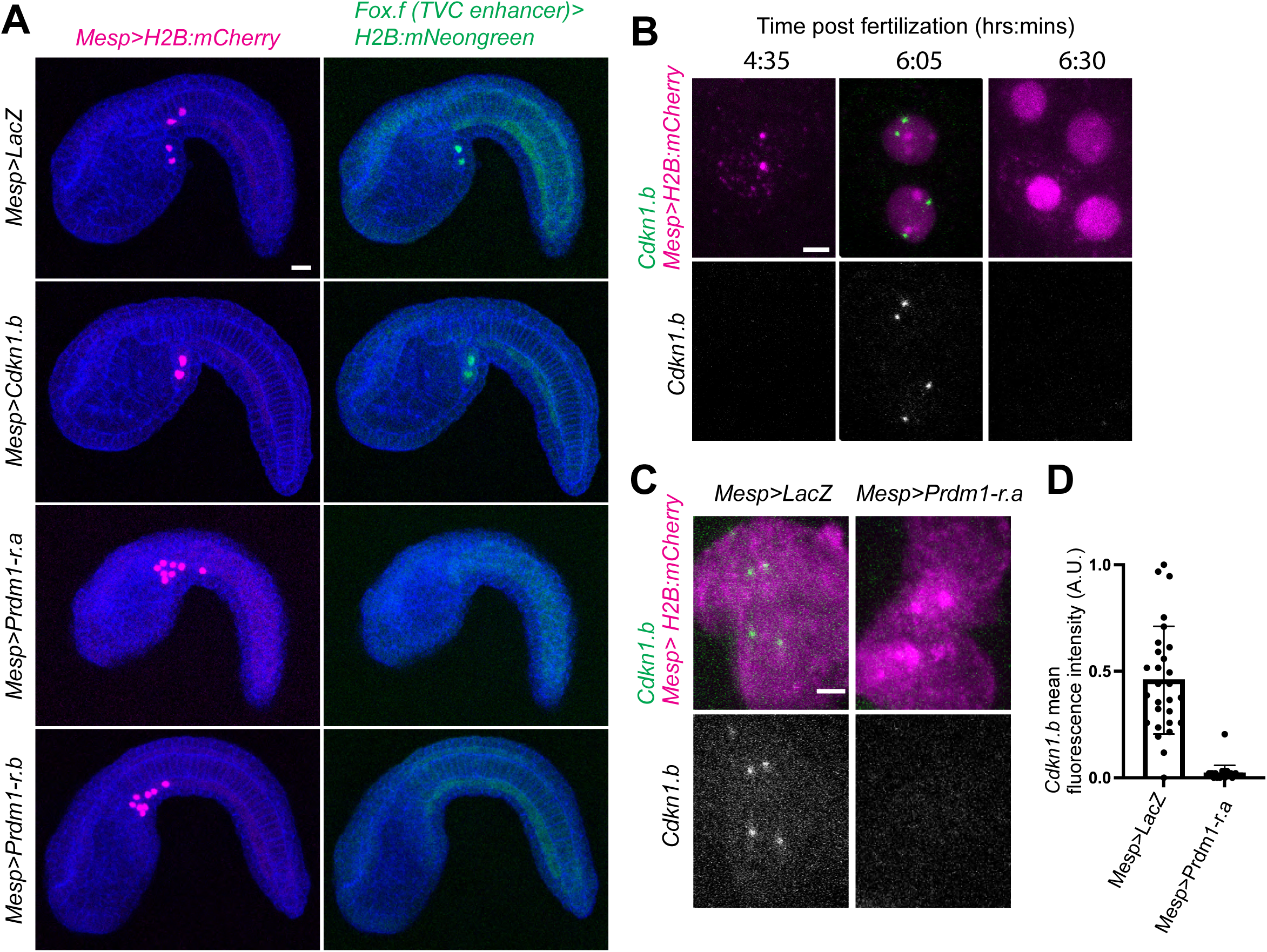
B7.5 cell numbers are regulated by *Prdm1-r.a/b* and *Cdkn1.b.* (A)Tailbud stage *Ciona* embryos electroporated with the indicated plasmids. Embryos have been stained with phalloidin (in blue) to visualize cell outlines. Cells that show red fluorescent signals in the nucleus (from H2B::mCherry) can be assumed to also harbor the other electroporated plasmids. Scale bar = 20 μm. (B) Expression of *Cdkn1.b* in B7.5 cells. B7.5 cells are labeled in magenta for fluorescent signals for endogenous *Mesp* and/or an H2B::mCherry reporter. HCR *in-situ* hybridization signals for *Cdkn1.b* (green or white) can only be seen at the 6:05 timepoint. Scale bar = 5 μm. (C) Expression of *Cdkn1.b* in control embryos and embryos where *Prdm1-r.a* is overexpressed. B7.5 cells are labelled in magenta for expression of the *mCherry* RNA, *Cdkn1.b* RNA is shown in green and white. Scale bar = 5 μm. (D) Quantifications of *Cdkn1.b* transcription in control (*Mesp>LacZ*) and *Prdm1-r.a* overexpressions (*Mesp>Prdm1-r.a*). Graph bars show means, error bars show standard deviations, and dots show individual measurements. The two conditions are significantly different when assessed by a two-tailed t-test (P<0.0001).

The cell cycle inhibitor *Cdkn1.b* has an important role in early ascidian embryos to stop mitosis at specific times in development allowing cell lineages to acquire precise cell numbers needed for correct development (20,21,24). We investigated the expression of *Cdkn1.b* in the B7.5 lineage and found no detectable *Cdkn1.b* RNA signal at the 110-cell stage, but clear signals for active transcription could be detected in daughter, B8.9-10 cells (Fig. 3B). This transcriptional signal was no longer detectable after the next mitosis in the B9.17-20 cells. Presumably this pulse of *Cdkn1.b* transcription is responsible for the arrest of cell division in B9.17-20 cells.

*Cdkn1.b* transcription was almost completely undetectable upon *Prdm1-r.a* overexpression in the B7.5 cell lineage (Fig. 3 C,D). We conclude that the increase in cell numbers upon *Prdm1-r.a* overexpression is due to repression of *Cdkn1.b* transcription, resulting in continued proliferation.

### Altering B7.5 lineage cell-numbers results in extra tail muscles

Because cell number was altered in *Cdkn1.b/Prdm-r* overexpression, we investigated if cell fate was changed. The B7.5 cells originate from the posterior blastomeres that inherit the muscle determinant *Macho1* (25) and two of the B7.5 granddaughter cells form anterior tail muscles. Therefore, the B7.5 lineage appears poised to form tail muscle. We looked for the expression of tail muscle markers upon overexpression of *Cdkn1.b/Prdm-r.a* (Fig. 4). The histone methyltransferase *Smyd1* and the muscle myosin component *Myl.g* are both highly expressed in the tail muscles and not in the TVCs (26,27). In a control electroporation (*Mesp>LacZ*, Fig. 4A) the anterior most cells that have begun migrating have markedly lower levels of *Smyd1* and *Myl.g* RNAs compared with posterior cells that form tail muscles. Surprisingly, when *Cdkn1.b* is overexpressed, both B8.9-10 cells exhibit high levels of *Smyd1* and *Myl.g*. When *Prdm1-r.a* or *Prdm1-r.b* are overexpressed the levels of *Smyd1* and *Myl.g* are mostly reduced (Fig 4.A S5). These results suggest that one or more myogenic determinants are asymmetrically segregated in TVCs and anterior tail muscles during the second division of the B7.5 lineage (see discussion).

**Fig. 4.**
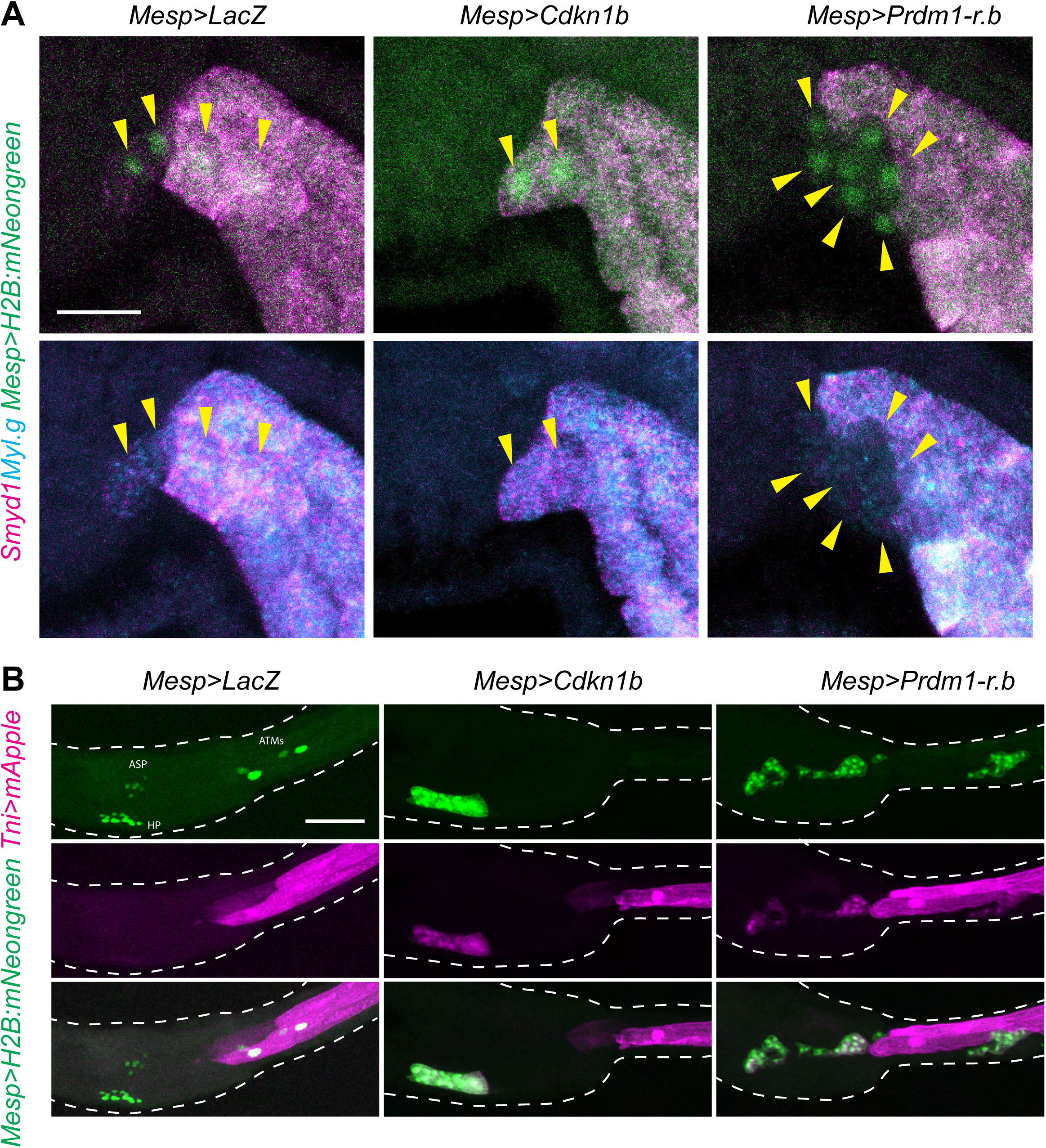
Disruption of B7.5 cell numbers disrupts cell fate. (A) HCR *in-situ* hybridization signals for *Smyd1* and *Myl.g* in tailbud stage *Ciona* embryos electroporated with the indicated plasmids. Cells with nuclei showing green nuclear fluorescence from the *Mesp>H2B::mNeongreen* plasmid can be assumed to harbor the other plasmids and are indicated by yellow arrowheads. Embryos are oriented anterior left, dorsal up. Scale bar = 20 μm. (B) Expression of muscle cell reporter (*Tni>mApple*) in *Ciona* larvae electroporated with the indicated plasmids. Larvae outlines are indicated by white dotted lines. ASP - atrial siphon primordia, ATMs – anterior tail muscles, HP – heart primordia. Larvae are oriented anterior left, dorsal up. Scale bar = 50 μm.

Because these overexpression assays are restricted to the B7.5 lineage, we obtained reliable development to larval stages to see the consequences of disrupting B7.5 cell numbers. In control experiments, each B7.5 blastomere forms two anterior tail muscles that express a *Tni* (troponin-1) muscle reporter plasmid (28). They also produce two migrating TVCs that will go on to form atrial siphon muscles and cardiomyocytes (Fig. 4B 4). When *Cdkn1.b* is overexpressed, the entire B7.5 lineage migrates into the trunk region, proliferates, and appear to form normal TVCs. However, more careful inspection reveals high levels of Tni reporter gene expression and a morphology consistent with small tail muscle cells (Fig, 4B). These observations suggest an uncoupling of TVC migration and specification, and the transformation of TVCs into supernumerary anterior tail muscles.

Overexpression of *Prdm1-r.a* and *Prdm1-r.b* results in more variable phenotypes. Some cells could be found in the trunk region, although these lack *Fox.f* expression and did not appear to have fully migrated. These cells sometimes formed loose and extended chain-like patterns with variable expression of the *Tni* reporter. We conclude that these cells have some of the properties of undefined progenitor cells, although a few of the cells appear to acquire a muscle-like identity.

## Discussion

The formation of ascidian tail muscles by unequal segregation of maternal determinants is one of the earliest known examples of cell lineage specification (29,30). This also remains one of the clearest understood embryonic processes with a complete causal link from maternal inputs to the expression of genes that trigger muscle differentiation and elongation (31). However not all cells that inherit the maternal determinants form muscle cells. For example, Macho1 is required for B5.1 cell descendants to form muscle, but cells from this lineage also form endoderm or notochord (31). This would suggest that muscle fates can be overridden by cell-signaling cues such as Nodal (32).

Previous studies on the specification of tail muscles and cardiopharyngeal mesoderm from the B7.5 lineage were interpreted in the context of this type of “signal override” mechanism. Due to its inheritance of maternal factors, the B7.5 lineage is fated to form muscle cells. At the early neurula stage, a localized FGF signal is received by the anterior B7.5 granddaughter cells, which overrides the muscle developmental program to trigger TVC migration and formation of atrial siphon muscles and cardiomyocytes (7).

Our observation that cell cycle arrest of B8.9-10 cells results in the migration of supernumerary anterior tail muscles suggests that the signal override model is not sufficient to account for the specification of the TVCs and cardiopharyngeal mesoderm.

Our results complement previous observations that inhibiting the function of *Fox.f* by overexpressing a modified, repressive form of the protein stops TVCs from migrating, but they are still able to develop into an ectopically positioned heart (22). By arresting cell division in the B7.5 lineage, we have produced the inverse outcome, migrating cells that only form tail muscle.

In addition to FGF signaling we suggest that specification depends on the asymmetric segregation of one or more muscle determinants during the second division cycle of the B7.5 lineage. Maternal *Macho1* mRNAs are undetectable in B7.5 cells (14,16) so it is an unlikely candidate for asymmetric segregation. However, Macho1 proteins can persist after mitosis (33) and it is possible that its degradation (along with other muscle determinants) is tightly coordinated with the cell cycle. The failure to divide beyond the B8.9-10 stage could result in the persistence of muscle determinants, despite activation of the *Fox.f* migratory program. Chromatin modifying transcription factors such as the histone methyltransferase Smyd1 could function as part of a positive feedback process that maintains myogenesis and blocks alternate fates

Another, possibly complementary, explanation could be that the muscle determining factors such as macho-1 remain localized in an anterior region of the B8.9 and B8.10 cytoplasm and these cells subsequently undergo an unequal division to deposit myogenic factors in the cells forming tail muscle (Fig. 5). If cell division is stopped early, these factors will remain and allow the cells to develop into muscle after migration has finished. The large amount of macho1 protein remaining in the posterior cytoplasm of the B5.2 cells at the 16-cell stage supports this interpretation (33).

**Fig. 5.**
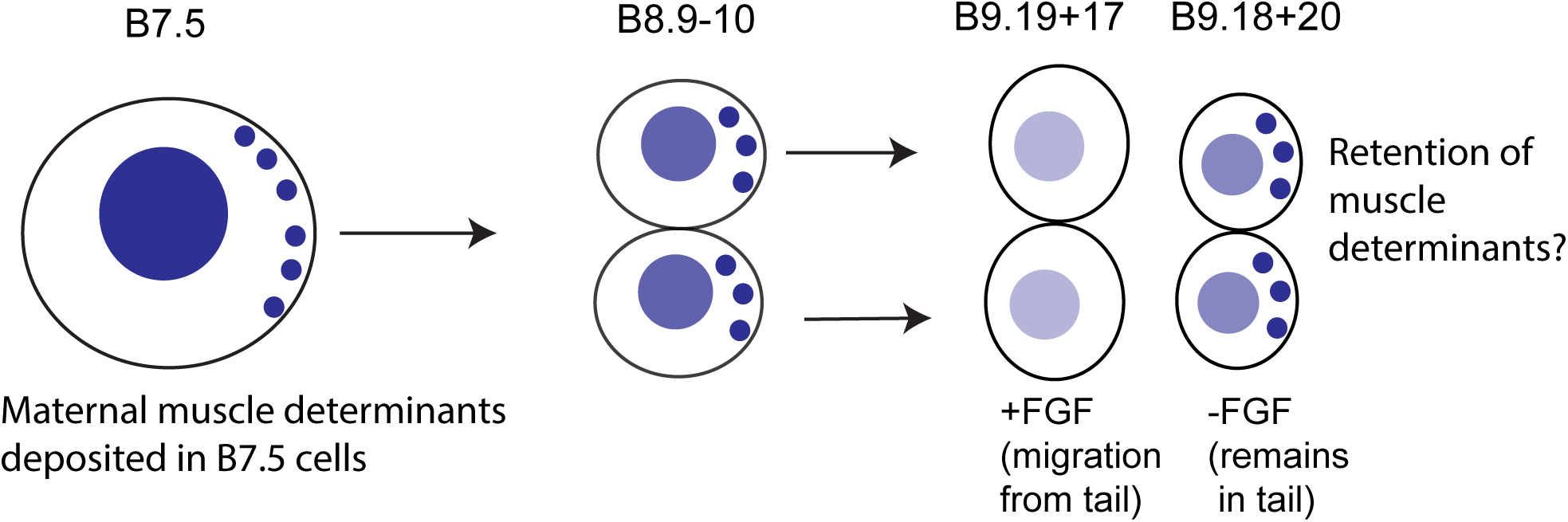
The connections between establishment correct cell number and correct cell fate in *Ciona* cardiopharyngeal mesoderm specification. A schematic showing a single B7.5 cell dividing twice to produce 4 granddaughter cells. The blue color represents the muscle forming factors present within the B7.5 cell and descendants that we propose is being diluted, degraded, or asymmetrically partitioned after the second mitosis. At the B8.9-10 stage the muscle forming factors are still present in a high enough concentration that they could activate muscle differentiation even if the cells receive an FGF signal to specify TVCs.

This study has highlighted the importance of cell numbers in embryonic development, and how changes in cell numbers can cause transformations in cell identity. Congenital heart defects are widespread, occurring in around 1% of births (34). In mice, Prdm1 mutants have reduced proliferation in the second heart field (35), suggesting a role of Prdm1 in regulating the numbers of mammalian heart progenitor cells. It is possible that these reduced numbers might also contribute to transformations in cell identity as seen in *Ciona*.

## Materials and Methods

### Single cell transcriptomic profile of 110-cell stage embryos

A previously published single-cell RNA-seq database of gene expression (14) was used to generate gene expression profiles using the tools on the Broad Institute Single Cell Portal (https://singlecell.broadinstitute.org). Genes were chosen based on previous literature describing their expression in early embryos.

### Animals

Adult *Ciona intestinalis* (pacific populations, type-A, also referred to as *Ciona robusta*) were supplied by M-REP (San Diego, CA) and Marinus Scientific (Long Beach, CA). Animals were stored in aerated aquariums filled with artificial sea water. Gametes were harvested surgically from adult *Ciona* and all procedures using live embryos were performed at 18°C.

### HCR *in situ* hybridization

Hybridization chain reaction (HCR) *in situ* hybridization was done as previously described (18). HCR probes and amplification reagents were produced by Molecular Instruments (Los Angeles, CA). Probe and hairpin combinations are described in S1.

### Quantification of transcription

Levels of transcription were measured as previously described (18). In brief, the spot within a nucleus with the highest fluorescent signal was assumed to be a site of active transcription. The mean fluorescence intensity of these spots was measured using the Zeiss Zen Blue software (Version 2.3, Zeiss, Oberkochen, Germany). If no bright spot was detected, a random spot within the nucleus was measured. Two measurements were made for each nucleus. Fluorescence intensity was normalized for each gene by subtracting the value of the lowest measurement for all measurements and then dividing by the value of the highest measurement. This produced a gene-specific normalized value of 0-1 for each measurement.

### Plasmid construction

The following novel plasmids were constructed for this study: *Mesp>LacZ*, *Mesp>Cdkn1.b*, *Mesp>Prdm1-r.a*, *Mesp>Prdm1-r.b*, *Mesp>H2B::mCherry*, *Mesp>H2B::mNeongreen*, *FoxF(TVC enhancer)-Fog(minimal promoter)>H2B::mNeongreen, Tni>mApple*. In all cases this nomenclature refers to the ‘regulatory region’>‘protein coding cDNA’. The backbone for all plasmids was a pSP plasmid (36). Genomic coordinates for regulatory regions and PCR Primers used to amplify DNA fragments for plasmid constructions are listed in Table S2. Regulatory regions were amplified from *Ciona* genomic DNA. *Ciona* protein coding gene cDNA were amplified from mixed embryonic stage cDNA using Primestar Max DNA polymerase (Ver 2, Takara Bio) according to the manufacturer’s instructions. Oligonucleotide primers used were the sequences described in Table S2 along with a 20bp 5’ sequence that overlapped the fusion target. Agarose gel extracted, purified PCR products were fused to form circular plasmids using an NEBuilder HiFi DNA Assembly Master Mix (New England Biolabs) according to the manufacturer’s instructions. Plasmids sequences were confirmed by whole plasmid sequencing (Plasmidsaurus, Louisville, KY). Full sequences are available upon request.

### Electroporation

1-cell stage *Ciona* zygotes were electroporated between 15-25 minutes post fertilization in 0.77M mannitol-artificial seawater as previously described (37). 20 µg of each plasmid was electroporated in an 800 µL electroporation with the exception of *Fox.F(TVC enhancer),Fog(minimal promoter)*>*H2B::mNeongreen* where 60 µg was electroporated.

### Microscopy

Imaging was performed using a Zeiss 880 confocal microscope. For HCR *in-situ* hybridizations, imaging conditions, including emission filter settings are the same as previously described (18). For imaging of reporter plasmids in larval stages, larvae were fixed in 2% formaldehyde, 0.5M NaCl, 0.1M MOPS, 1 mM EGTA, 2mM MgSO4, 0.1% Tween-20 for 30 minutes and washed 4 times in PBST. Embryos were then imaged using a Zeiss 880 confocal microscope. Additional specifics on imaging conditions are available upon request.

## Acknowledgements

We thank members of the Levine lab for helpful discussions. This work was supported by funds from the Princeton Catalysis Initiative (PCI).

## Supporting Information Figure Legends

**Fig. S1.**
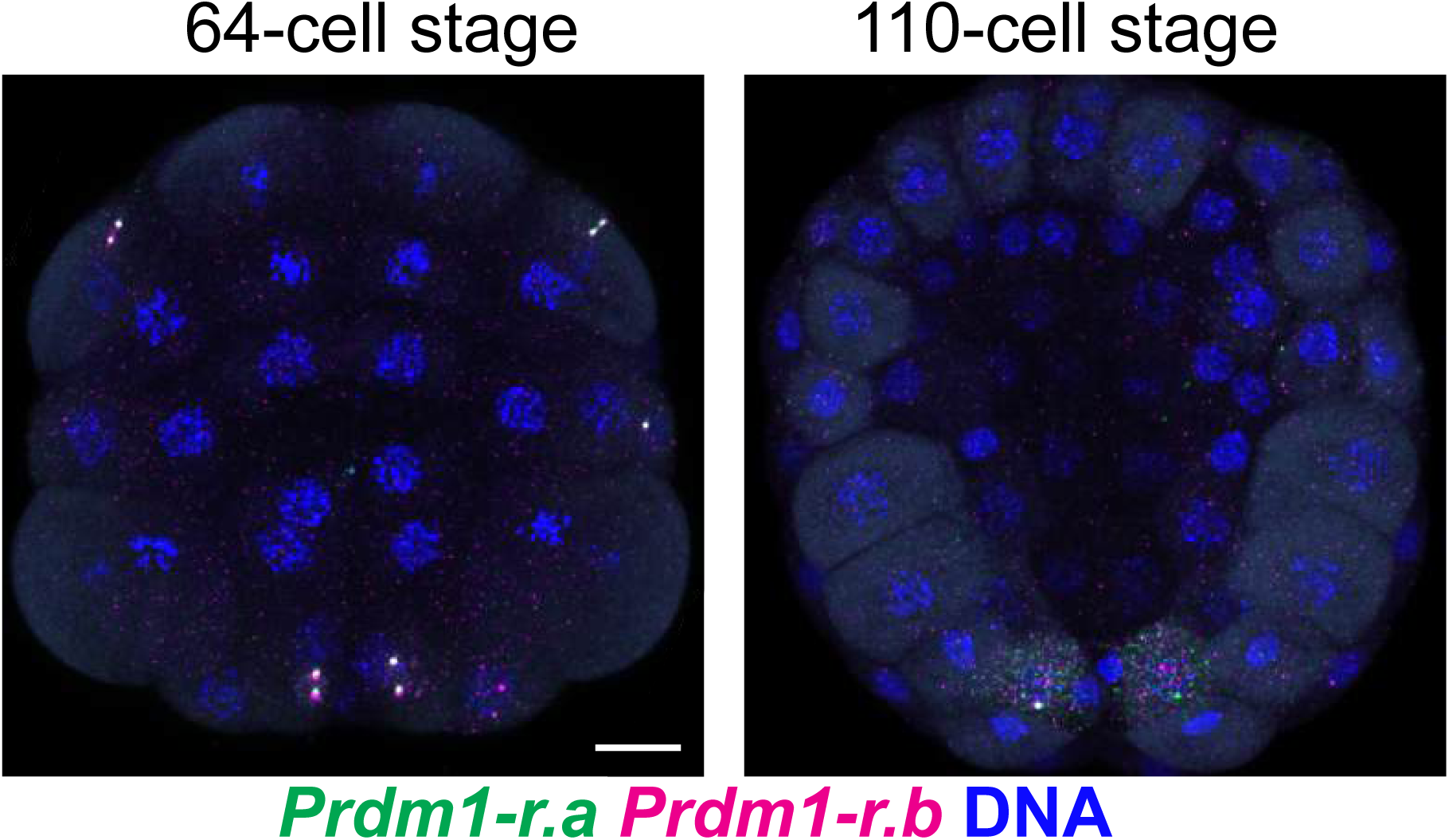
*Prdm1-r.a* and *Prdm1-r.b* expression at the 64- and 110-cell stages. HCR *in-situ* hybridization signals for *Prdm1-r.a* and *Prdm1-r.b*. Images are extended maximum intensity projections of the same dataset depicted in Fig. 1C. Embryos are vegetal views, anterior up. Scale bar = 20 μm.

**Fig. S2.**
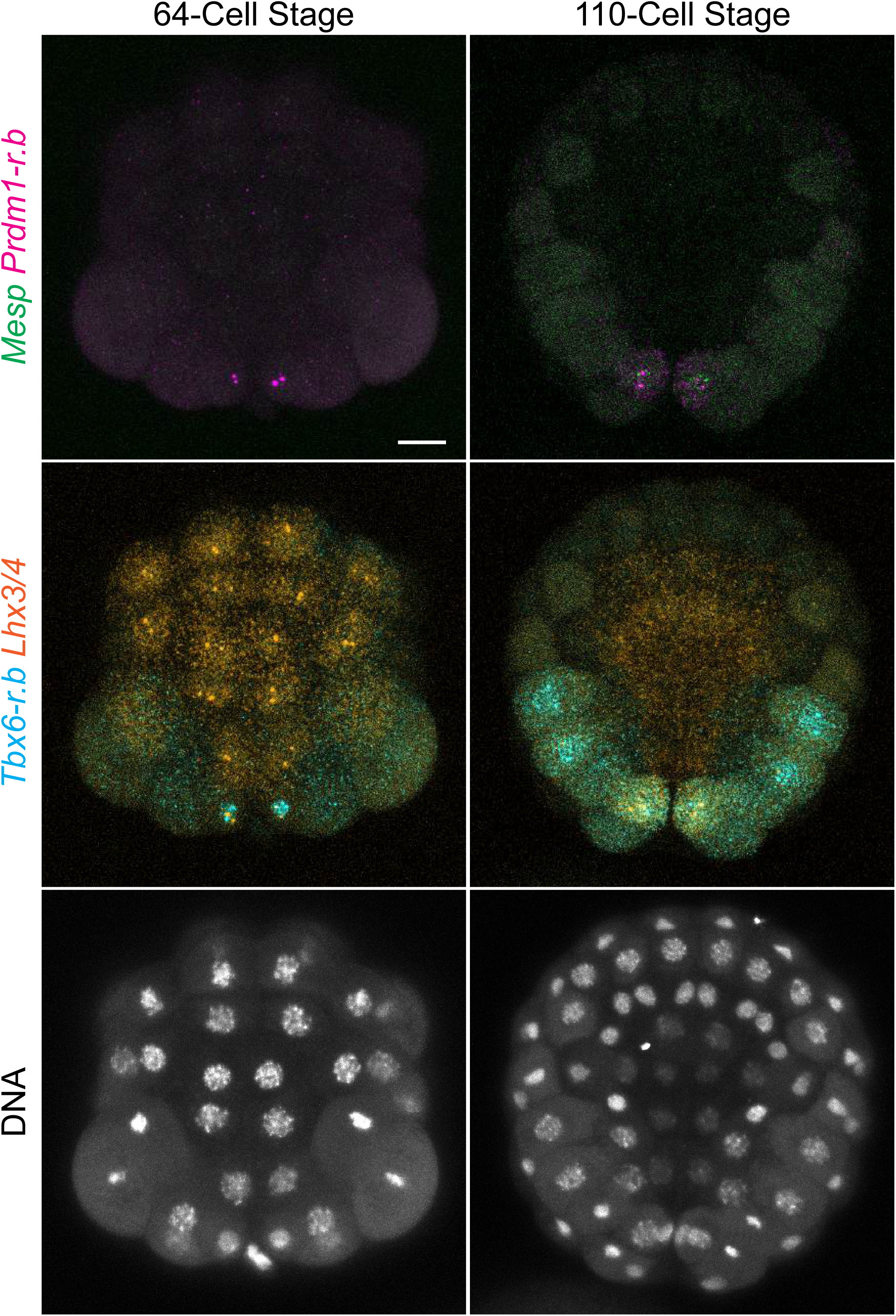
4-color HCR *in situ* hybridization in 64- and 110-cell stage *Ciona* embryos. HCR *in-situ* hybridization signals for *Tbx6.b, Lhx3/4*, *Prdm1-r.b*, and *Mesp* in Ciona embryos at the 64- and 110-cell stages. The 3 images in a single time point are from the same embryo. Images are whole embryo views of samples from the same dataset shown in Fig. 2. Embryos are vegetal views, anterior up. Scale bar = 20 μm.

**Fig. S3.**
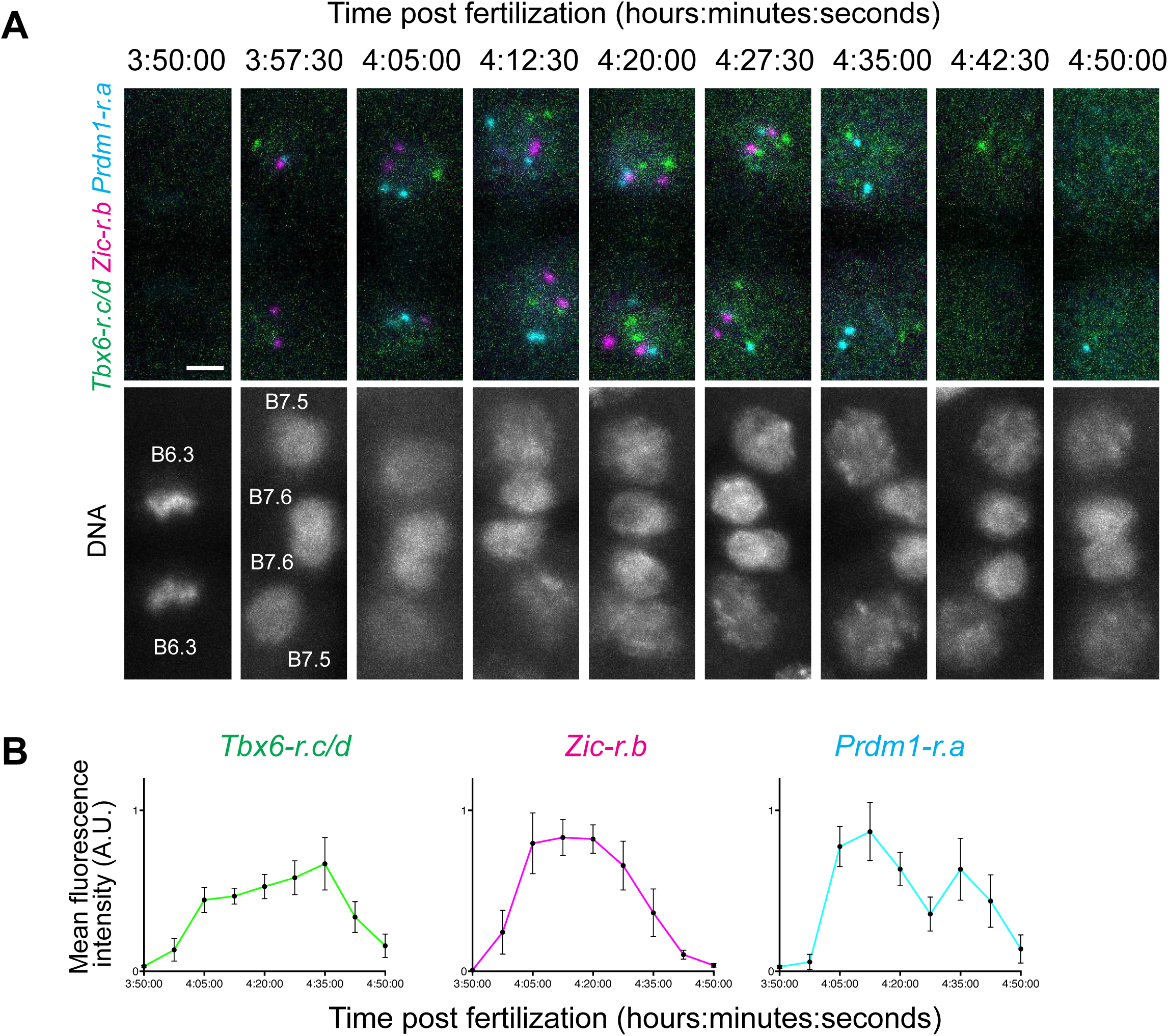
Additional measurements of onset of transcription in B7.5 Cells. (A) HCR *in situ* hybridization signals for *Tbx6-r.c/d, Zic-r.b*, and *Prdm1-r.a* from the onset of mitosis of the B6.3 cells for 1 hour. Images are taken from individual fixed samples. The 3 images in a single time point are from the same embryo. Scale bar = 5 μm. (B) Quantification of levels of transcription from HCR *in situ* hybridization signals for *Tbx6-r.c/d*, *Zic-r.b*, and *Prdm1-r.a.* Each time point consists of 28 measurements from 7 embryos. Error bars plot 95% confidence intervals.

**Fig. S4.**
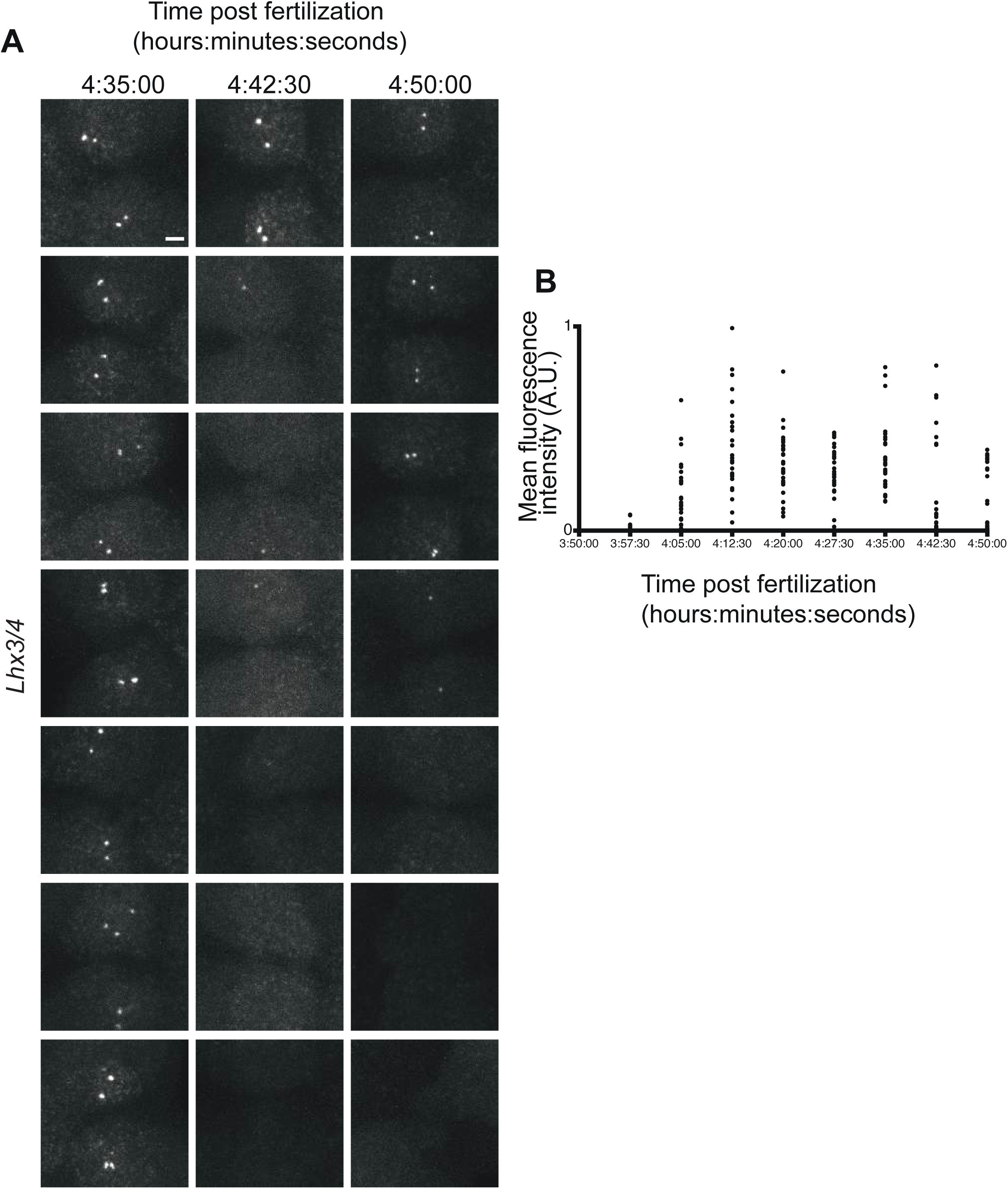
Lhx3/4 exhibits behavior consistent with transcriptional bursting. (A) HCR *in situ* hybridization signals for *Lhx3/4* in B7.5 cells. Each image is from an individual embryo. Scale bar = 5 μm. (B) Quantification of levels of transcription from HCR *in situ* hybridization signals for *Lhx3/4* from the images in A. This figure uses the same dataset as Fig. 2. but depicts the individual measurements.

**Fig. S5.**
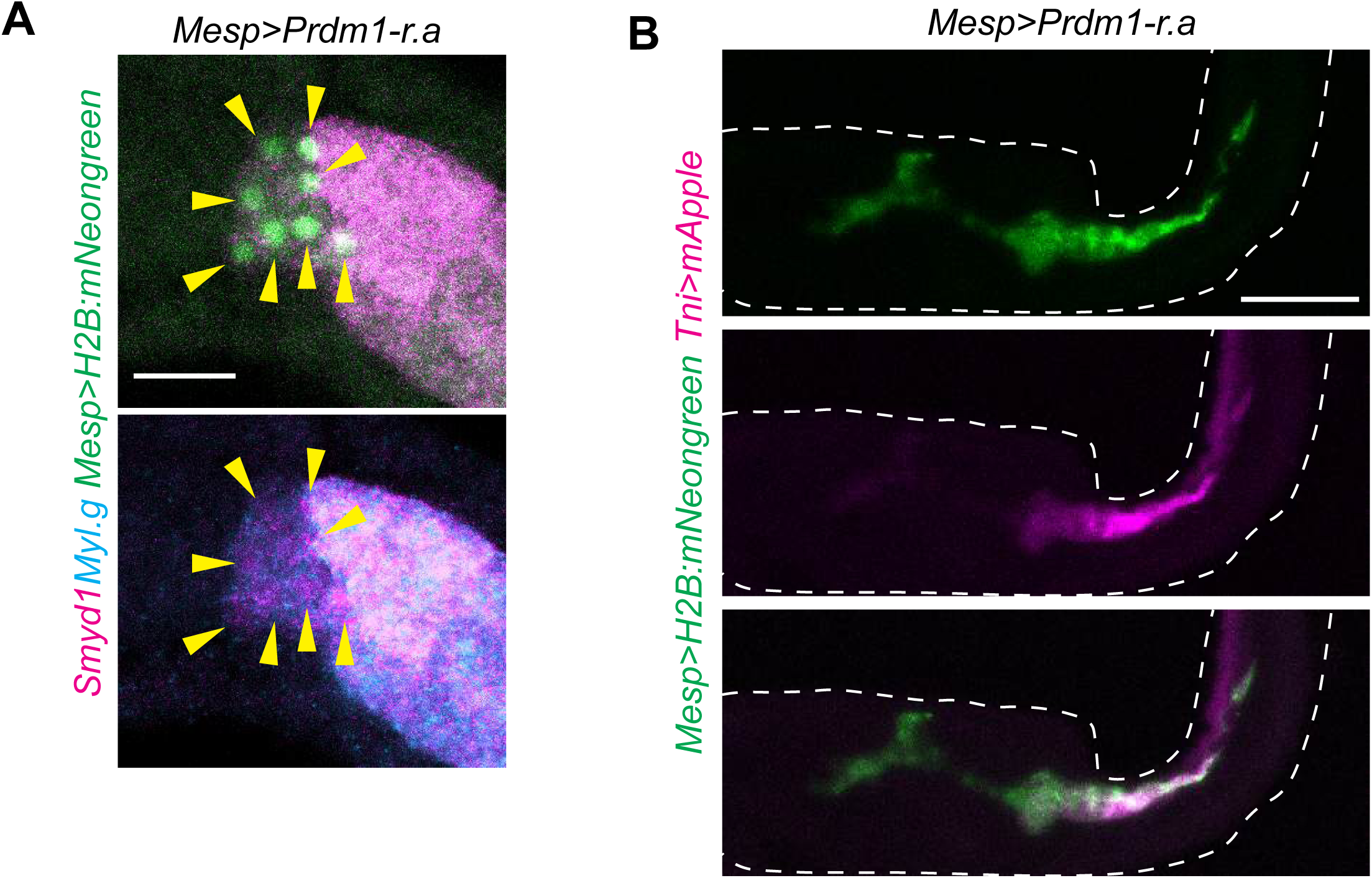
Additional evidence disruption of B7.5 cell numbers disrupts cell fate. (A) HCR *in situ* hybridizations and fluorescent reporter experiments showing the same data in Fig. 4 but for *Prdm1-r.a*. Expression of muscle genes *Smyd1* and *Myl.g* in a tailbud stage *Ciona* embryo electroporated with the indicated plasmids. Cells with nuclei showing green nuclear fluorescence from the *Mesp>H2B::mNeongreen* plasmid can be assumed to harbor the other plasmids and are indicated by yellow arrowheads. Embryo is oriented anterior left, dorsal up. Scale bar = 20 μm. (B) Expression of muscle cell reporter (*Tni>mApple*) in *Ciona* larva electroporated with the indicated plasmid. Larva outlines are indicated by white dotted lines. Larva is oriented anterior left, dorsal up. Scale bar = 50 μm.

**Table S1.**
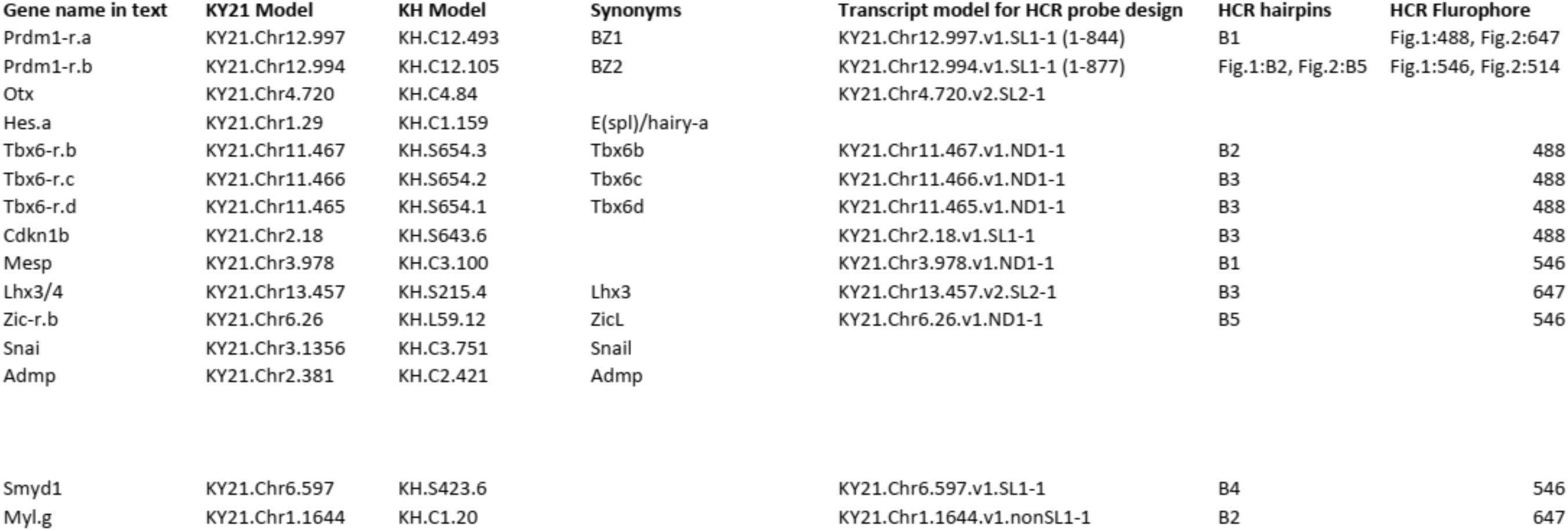

